# Neurotranscriptomic signatures of natural variation in mate preference learning in two subspecies of *Heliconius melpomene* butterflies

**DOI:** 10.64898/2026.02.15.706038

**Authors:** Sushant Potdar, Kiana Kasmaii, Chance Powell, Erica L. Westerman

## Abstract

Many animals change their behavior in response to social experiences by learning. Although social learning is adaptive, not all individuals learn. In *Heliconius melpomene, H. m. malleti* males respond to 2-day prior failed copulation experience by decreasing courtship, whereas *H. m. rosina* males do not. Here, we explore the transcriptomic differences in both the neural (brain) and sensory (eyes, antennae) tissues underlying this natural diversity in male aversive mate-preference learning. While the transcriptomic profiles of the two subspecies are inherently different across all three tissues, we found the greatest difference between the good (*H. m. malleti*), and bad (*H. m. rosina*) mate-preference learners in the brain, followed by the sensory tissues. Known learning genes and Gene Ontology terms were associated with differences in mate-preference learning, suggesting conserved learning pathways across animals. Genes within putative magic loci associated with colors, odors, and locomotion, were also differentially expressed between *H. m. malleti* and *H. m. rosina,* suggesting multimodal sensory processing may drive behavioral variance in these two subspecies. Overall, our study identifies genetic underpinnings for differences in preference learning, both in neural processing and sensory tissues. Selection on these genes/networks could result in preference learning-induced reinforcement, leading to reproductive isolation and speciation.

## Introduction

Learning, where individuals change their behavior by incorporating past experiences, is prevalent throughout the animal kingdom, from cnidarians to chordates^1^. Learning can be adaptive^2,3^, but not all animals learn. Even within the same species, differences in local selection pressures, social contexts, or the developmental environment can result in variance in learning ability^2–4^. Understanding the molecular mechanisms underlying variation in learning ability informs how developmental environment and evolutionary forces maintain variation in this important adaptive trait^5^.

One common type of learning is mate preference learning, when individuals use past social experiences to form or change their mate preferences^6–9^. Mate preference learning can be adaptive, as it fine tunes preferences based on past experience, and prevents courtship and mate choices that reduce fitness^7,10,11^. But learning can also be costly, requiring time and energy and increasing the risk of maladaptive preferences^2,3,12,13^. From a molecular perspective, selection should favor differential regulation of learning and memory associated genes under divergent selection for preference learning.

*Heliconius melpomene* are a strong model for exploring the transcriptomic mechanisms underlying variance in mate preference learning due to their morph diversity, relatively large brains, and documented learning ability^14–17^. *H. melpomene* learn color preferences associated with food rewards^16,17^, and also exhibit morph-specific variation in male mate preference learning^18^.

The genetic landscape of allopatric *H. melpomene* subspecies are similar, with a few genomic “hotspots” surrounding loci for wing color, pattern, and shape diverging due to natural and sexual selection^19–21^. Wing colors, patterns, shape, and sex-specific pheromones are used as sexual signals for assortative mating in *Heliconius*^22–29^. Genes controlling preference for wing color are physically linked and adjacent to loci regulating wing color and pattern development that are under divergent natural selection (“magic” traits)^30–33^. Wing color and patterning genes are implicated in sexually dimorphic imprinting-like learning in *Bicyclus anynana* butterflies^34^, but whether such pleiotropic genes that control the development of a “magic” trait (wing color/pattern/shape) and its preference also influence natural variation in learning ability has yet to be determined. Further, if preference genes, “magic” trait genes, and genes associated with differences in mate preference learning are physically linked (albeit not the same gene), then selection acting on one (eg, mating trait) could maintain the other two traits (eg. preference and learning) due to linkage. Identifying such pleiotropic loci provides a framework for understanding how selection on traits, preference, or learning can drive divergence through linkage.

Here, we explore the neurogenomic changes associated with differences in mate preference learning in two subspecies of *Heliconius melpomene* butterflies^18^. Brazilian and Ecuadorian *H. m. malleti* males decrease their courting of females after a previous (2-day prior) exposure with a female without mating, whereas Panamanian and Costa Rican *H. m. rosina* males do not^18^. Here we assess the neurogenomic causes of these differences in mate preference learning by examining differentially expressed genes (DEGs) in brain (BR), eye (EY), and antennae (AN) tissues between *H. m. malleti* and *H. m. rosina* males just after a social exposure with a female of the same phenotype (training treatment; Figure S1). We compare the transcriptomic signatures in the absence of a female (control) to establish baseline expression differences between subspecies (Figure S1). Next, we identify shared DEGs between our study and those differentially expressed in the brains of *H. melpomene* and *H. cydno* during male mate preference^18,31^, and in the developing wings of *H. melpomene*^37^. Lastly, we searched for differentially expressed putative “magic” genes, genes controlling known sexual signaling traits, and genes on the Z sex chromosome. We postulate that learning and memory genes may be linked with “magic trait” loci. We also hypothesize that, while *H. m. rosina* have the ability to learn spatial cues for floral rewards^16,17^, learning genes may not be expressed and/or disassociated in a mate preference context, which may contribute to the observed similarity in courtship frequency in trained and naïve *H. m. rosina* males^18^, irrespective of social experiences.

## Results

### H. m. malleti and H. m. rosina have baseline transcriptomic differences in brain and sensory tissues

We first determined whether *H. m. malleti* and *H. m. rosina* males had inherent differences in the brain and the sensory tissues by comparing gene expression of control (isolated free flying) 10-day old males (Figure S1 A, B). We found over a thousand DEGs between subspecies for each tissue (Figure 1A), with1608 DEGs in the brain (Table S2, Figure S3E), 1301 DEGs in the eyes (Table S3, Figure S3C), and 1398 DEGs in the antennae (Table S4, Figure S3A). Of these, 62%-72% per tissue were uniquely DE in the control setting, and included genes associated with neural development and signaling, sexual reproduction, and hormones (Figure 1B-E, Tables S5, S6, S7). Of these uniquely DEGs, 54 were common between all three tissues, including two genes we found particularly intriguing: *Nardilysin* (HMEL008206g5), which is involved in axonogenesis; and a ligand transporter apolipoprotein (HMEL030910g1; Figure S4A, Table S16) involved in brain development and peripheral axon regeneration. This large set of DEGs in free flying and isolated males suggest that the two populations may have different sensory umwelts, and inherent differences in sensory and neural processing (for more details see Supplementary results).

**Figure 1:**
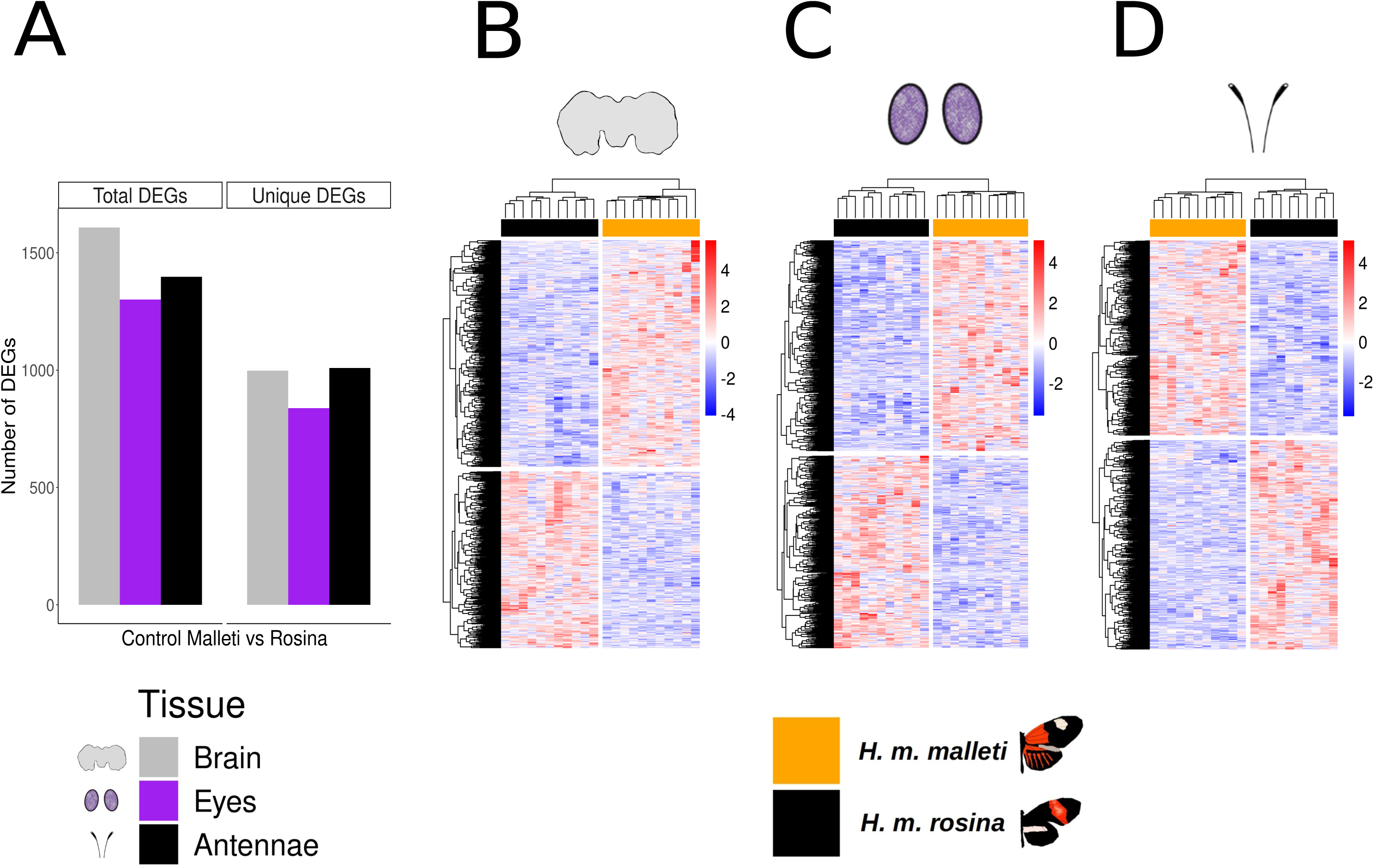
*H. m. malleti* and *H. m. rosina* males are transcriptomically different. A) Transcriptomics analysis revealed similar number of differentially expressed genes (DEGs) between 10-day-old sexually mature males of *H. m. malleti* and *H. m. rosina* in the brain, eyes, and the antennae tissues. Roughly 62-72% of DEGs across the three tissues were uniquely differentially expressed in the control conditions only. (B-D) Heat maps of uniquely control condition specific DEGs segregate males of *H. m. malleti* and *H. m. rosina* separately in the (B) brain, (C) eyes, and (D) antennae tissues. In these heat maps, each row is a single gene, and each column is an individual sample (indicated by cluster trees in both rows and columns). Gene expressions are plotted as Z-scores after variance-stabilizing transformation, with red colors indicating increased expression, and blue colors indicating decreased expression relative to the mean for that gene.

### Variation in learned response is associated with more DEGs in the brain than in sensory tissues

To assess transcriptomic variation associated with differential mate preference learning in *H. melpomene* (Figure 2A), we performed RNAseq and compared gene expression in the same three tissues (BR, EY, AN) of *H. m. malleti* and *H. m. rosina* males collected immediately following a 90 minute exposure to a female of their own phenotype (training treatment, Figure S1 C, D). We recovered 1929 DEGs in the brain (Table S9, Figure 2A, Figure S3F), 1201 DEGs in the eye (Table S10, Figure 2A, Figure S3D), and 834 DEGs in the antennae (Table S11, Figure 2A, Figure S3B). Of these, 53-68% were unique DEGs (Table S12-S14, Figure 2), and are not DE between control *H. m. malleti* and *H. m. rosina* (see above). Many of these DEGs are associated with learning and memory, sexual reproduction, neural signaling and neurodevelopment, and zinc ion binding categories, among others (Figure 3B-D). Of the uniquely DEGs in the training contrast, 29 were common between all three tissues (Table S17, Figure S4B). Of particular note are a neural signaling and neurodevelopment gene *Potassium voltage-gated channel protein Shab* (*Shab*, HMEL022148), which was downregulated in trained *H. m. malleti* in all three tissues (Figure S4D), *regucalcin* (HMEL034199g1), which was upregulated in the sensory tissues (EY, AN), but downregulated in the brain of *H. m. malleti* relative to *H. m. rosina*, and *HD domain-containing protein 2* (HMEL008530g1), which was upregulated in the brain and the eyes, but downregulated in the antennae of *H. m. malleti* relative to *H. m. rosina* (Figure S4D). These unique DEGs may be associated with the differential mate preference learning behavior of these two populations (Figure 2A).

**Figure 2:**
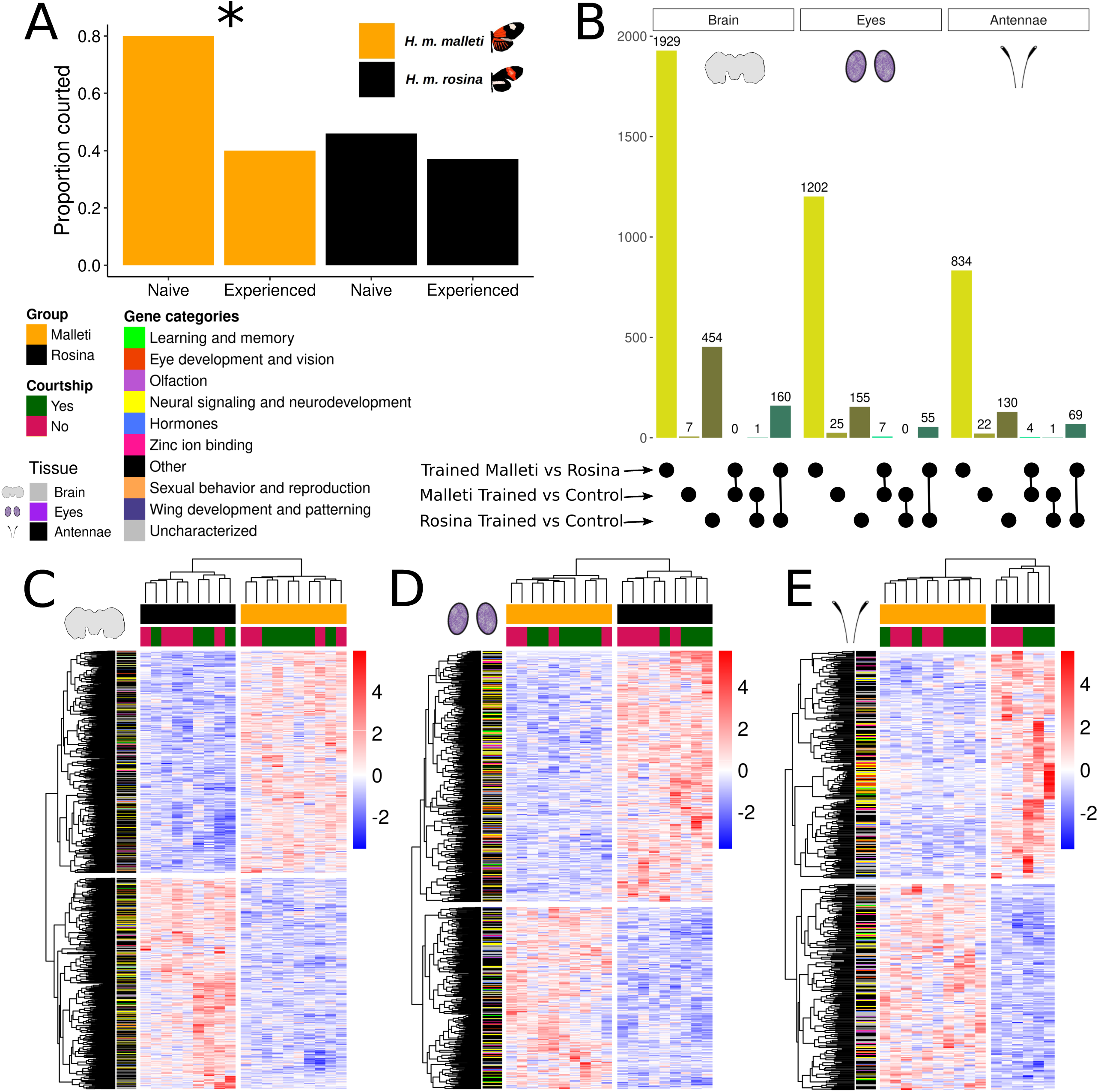
Greater DEGs in the brain is associated with the difference in mate preference learning between male *H. m. malleti* and *H. m. rosina*: A) Male *H. m. malleti* reduce courtship after a failed copulation experience (compared to naive), whereas *H. m. rosina* do not reduce courtship after a failed copulation experience ^18^. B) Brain tissue showed the highest number of DEGs between male *H. m. malleti* and *H. m. rosina* after a training experience with their respective female subspecies, followed by the eyes, and the antennae. Numbers on top of each bar plot indicates the number of DEGs. Dots on the x-axis indicates DEGs associated with that comparison, with dots joined by lines indicating common DEGs between those two comparisons. (C-E) Heat maps of uniquely training specific DEGs segregate male *H. m. malleti* and male *H. m. rosina* after training with a female in the (C) brain, (D) eyes, and (E) antennae tissues. DEGs are mainly clustered between subspecies, and sub-clustered between whether the males courted the females during a brief (90 minutes) interaction with them. Like in Figure 1, in these heatmaps, each row is a DEG, whereas each column is a sample. Warmer (red) colors indicate increased expression, and cooler (blue) colors indicate decreased expression compared to their means. These heat maps contain many gene categories that are important for processes associated with differences in mate preference learning between males of *H. m. malleti* and *H. m. rosina*.

**Figure 3:**
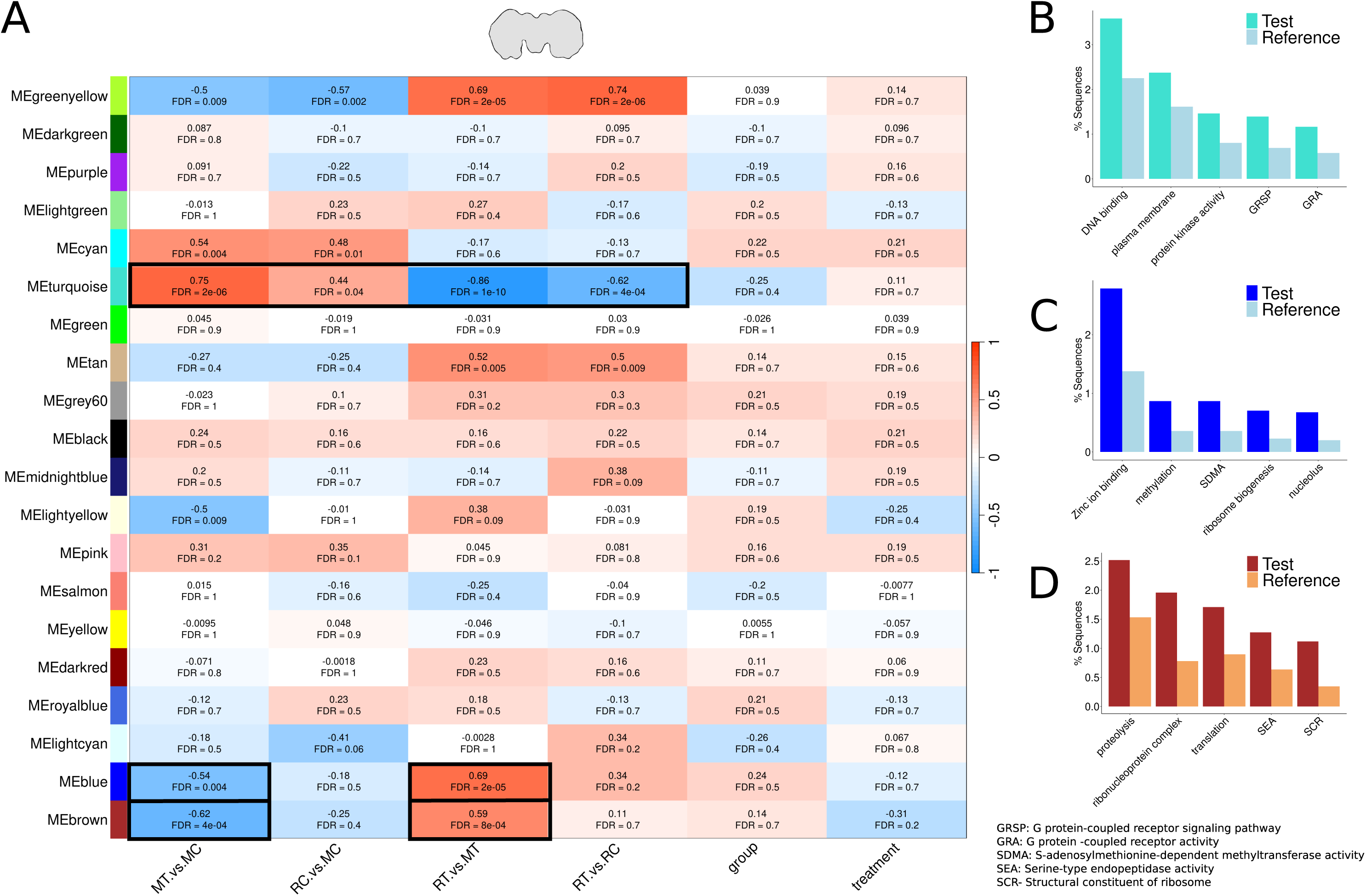
Two brain gene network modules are associated with differences in *H. melpomene* male mate preference learning: A) Heatmap of gene module-trait association indicates two modules (blue and brown) that are uniquely associated with female-associated differences between *H. m. malleti* and *H. m. rosina* (MT vs MC and RT vs MT), while the turquoise module is associated with the general transcriptomic differences between the two subspecies. In this heatmap, each row is a module eigengenes (ME), whereas each column is a trait comparison. In each box, the strength of association (from −1 to 1) and the false discovery rate (FDR) are reported. MT vs MC = *H. m. malleti* trained vs *H. m. malleti* control; RC vs MC = *H. m. rosina* control vs *H. m. malleti* control; RT vs MT = *H. m. rosina* trained vs *H. m. malleti* trained; RT vs RC = *H. m. rosina* trained vs *H. m. rosina* control; group = *H. m. malleti* vs *H. m. rosina*; treatment = trained vs control. (B-D) Many GO terms implicated in learning and memory formation were enriched in gene-network modules associated with differences in subspecies ((B) turquoise module), and associated with differences in mate preference learning ((C) blue, and (D) brown modules). In (B-D), the test set contains percentage of sequences annotated for that term among all the genes in the module, and the reference set contains the percentage of sequences annotated for that term in the transcriptome.

During training, 27 GO terms were significantly enriched and over-represented in *H. m. malleti* compared to *H. m. rosina* in the brain tissue (Table S15). Some terms that were enriched include “ATP binding”, and “glutathione transferase activity”, consistent with energetic demands needed for learning and memory^38^. Four glutathione S-transferase genes (HmelGSTs1, HmelGSTe4, HmelGSTd2, HmelGSTt1) were uniquely upregulated in trained *H. m. malleti* brains. Weighted Gene Co-expression Network Analysis (WGCNA) identified three modules in the brain (lightyellow, blue, brown; Table S24) that were significantly associated with differences between learning and non-learning contrasts (which includes trained vs control *H. m. malleti*; Figure 3A). GO enrichment identified 40 terms for the blue module (Figure 3C, Table S26) and 53 in the brown module (Figure 3D, Table S27). Most notable term in the blue module is “zinc ion binding” which regulates diverse neuronal processes from neurotransmission to signaling, learning, and memory formation (Figure 3C). In the eyes, WGCNA identified one module (yellow) that was significantly associated with the differences between *H. m. malleti* and *H. m. rosina* after an exposure with a female, however, this module is also associated with other contrasts (Figure S6A, Table S28). In the antennae, no significant GO terms nor WGCNA modules were associated with training induced differences between *H. m. malleti* and *H. m. rosina* antennae (Figure S6D, Table S31).

### Genes within wing color and patterning “magic” loci are DE between H. m. malleti and H. m. rosina

We next searched for genes/loci that control the wing color and patterning “magic” traits–which are under both natural and sexual selection^15,23,24^ –to identify whether they are DE between *H. m. malleti* and *H. m. rosina* in baseline and preference learning contexts. First, we examined the four known *Heliconius* wing color and patterning genes (*optix, cortex, WntA,* and *aristaless*), of which only *cortex* (HMEL000025, log2foldchange = −0.06) was DE between *H. m. malleti* and *H. m. rosina* (Figure 4B).

**Figure 4:**
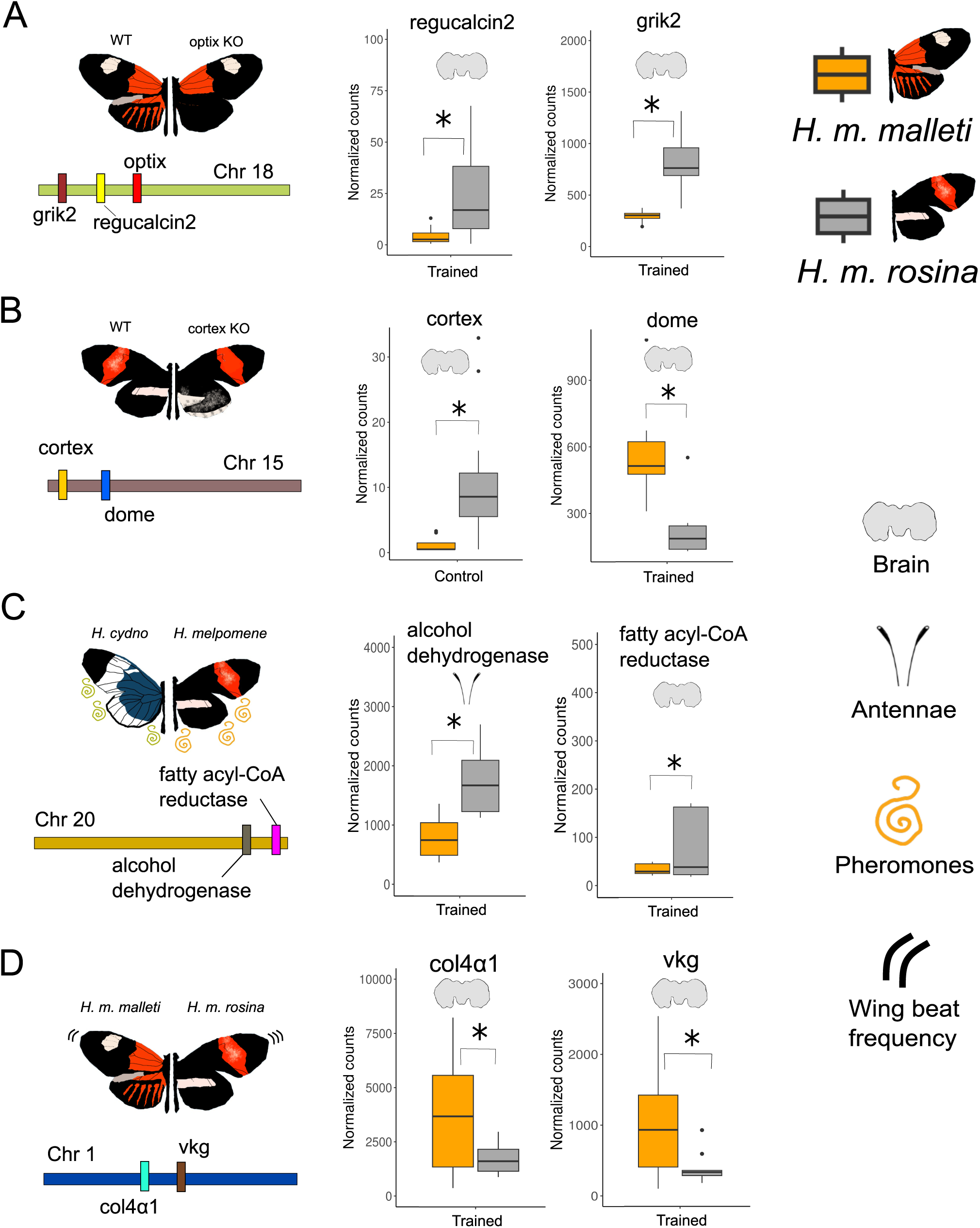
Putative magic loci genes may be implicated in mate preference learning: Pleiotropic genes/loci responsible for traits that are under both natural, and sexual selection, and their adjacently located genes are DE between *H. m. malleti* and *H. m. rosina*. These include A) Genes adjacent to the color patterning gene *optix,* that are important in visual mate preferences in *Heliconius* butterflies. B) Another *Heliconius* color patterning gene, *cortex*, and its adjacent gene *dome* and *wash* (not shown). C) Genes in the pheromone synthesis pathway, and D) Genes responsible for differences in wing beat frequency. In each panel (A-D), the function of the genes/loci are shown graphically, the genes’ location on the chromosomes (not to scale), and the differences in normalized counts of DE genes. *p<0.05.

Next, we looked at genes immediately adjacent to these four functionally characterized *Heliconius* wing patterning genes. Three genes downstream of the red ommochrome pigmentation gene *optix*^41,42^ are DE between *H. m. malleti* and *H. m. rosia*: *regucalcin 1* (*RGN-1*, HMEL013551g4), which is a preference gene for red wing color in *H. melpomene*^26^, is downregulated in *H. m. malleti* eyes; *regucalcin 2* (*RGN-2*, HMEL013552g1 and HMEL034199g1), which are uniquely DE during training between the two subspecies in all three tissues; and *Glutamate receptor ionitropic, kainate* (*GRIK2*, HMEL009992g4 and HMEL009992g1) is downregulated in *H. m. malleti* in all tissues (BR, EY, AN) and scenarios (trained and control; Figure 4A). While two genes adjoining *cortex–*which controls the wing scale identities upstream of color patterning genes *optix* and *aristaless1*^43^–are DE in the brain between *H. m. malleti* and *H. m. rosina*. The gene *wash* (HMEL000036, log2foldchange = 0.16) is upregulated in *H. m. malleti* brains in the control treatment while *domeless* (*dome*, HMEL013472g1) is upregulated in *H. m. malleti* brains in both control (log2foldchange = 1.43) and trained (log2foldchange = 1.26) treatments (Figure 4B; see supplementary results for *WntA* and *aristaless 1*).

We utilized previously published hindwing transcriptomic data to identify a suite of genes that are commonly DE between developing *H. m. rosina and H. m. melpomene* wings^37^ and male sensory and neural tissues (our study), suggesting an influence on both wing pattern and sensory perception or cognitive processing. We identified 40 DEGs in the antennae (Table S37); 101 DEGs in the eyes (Table S38); and 98 DEGs in the brain (Table S39) between *H. m. malleti* and *H. m. rosina* in the control treatment from our study that were common to the DEGs in the developing hindwings between *H. m. rosina* and *H. m. melpomene*^37^. Some important genes that were commonly differentially expressed in wing, and sensory and neural tissues include *regucalcin* (HMEL013552g1; see above), *Gamma-aminobutyric acid receptor subunit alpha-6* (*GABRA6,* HMEL007707g1) which is involved in regulating the inhibitory neurotransmitter GABA, and *protein eyes shut* (HMEL033043g1) which is required for retina formation^44^.

### Genes controlling sexual traits and preferences are DE between H. m. malleti and H. m. rosina

We also compared a suite of probable male mate preference genes that were differentially expressed between *H. m. malleti* and *H. m. rosina* brains in our study to brain DEGs between *H. m. rosina* and *H. cydno chioneus*^31^ (Table S40). We identified 114 genes differentially expressed in both data sets, including *GRIK2* (HMEL009992g1 and HMEL009992g4) that is upstream of the wing color gene *optix* (see above), a neurogenesis and zinc ion binding gene *shuttle craft* (HMEL013483g1), gene *yellow* (HMEL021221g1), and an *ommochrome binding protein* (HMEL030103g1). These genes may be important for visual mate preference in *Heliconius* butterflies on a broad scale, as they are differentially expressed between populations and species, and in butterflies that differ in wing pattern but not color (*H. m. malleti* and *H. m. rosina*) as well as butterflies that differ in wing color (*H. m. rosina* and *H. c. chioneus*).

Olfactory traits such as male sex pheromones are species specific in *Heliconius* butterflies^45^. Of the 26 candidate genes in the putative pheromone and anti-aphrodisiac biosynthesis pathway^46,47^, we identified 10 genes that were DE in our study (Table S41). One such gene is *geranylgeranyl pyrophosphate synthase* (*GGPS1*, HMEL037104g1) which is upregulated in *H. m. malleti* relative to *H. m. rosina* in brains, eyes, and antennae. Two other genes that were DE between *H. m. malleti* and *H. m. rosina* are *Fatty acyl-CoA reductase wat* (*wat*, HMEL014878g1) which is downregulated in *H. m. malleti* brain in the training treatment (log2foldchange = −1.82, Figure 4C) and a *D-arabinitol dehydrogenase 1* (*ARD1*, HMEL005376g1, log2foldchange = −0.82), which is uniquely downregulated in *H. m. malleti* brain relative to *H. m. rosina* (Figure 4C).

### Locomotion locus and wing shape genes are DE between H. m. malleti and H. m. rosina

Another locus that has consistently diverged across *Heliconius* is the L locus, which is involved in differences in locomotory behaviors^21^. Four genes at this locus were DE between *H. m. malleti* and *H. m. rosina* in this study. *Collagen α-1 (IV) chain* (*Col4α1,* HMEL030288g1) and *Collagen α-2 (IV) chain* (*Vkg,* HMEL002288g1) were upregulated in *H. m. malleti* trained brain (log2foldchange = 1.05, 1.44 respectively, Figure 4D), and *Sel-T like protein* (*SelT,* HMEL002290, log2foldchange = 0.47) was upregulated in *H. m. malleti* trained eyes. *WD repeat-containing protein 35* (*Oseg4,* HMEL002291g1) was also upregulated in *H. m. malleti* eyes during both the control (log2foldchange = 0.41) and the trained (log2foldchange = 0.6) treatment. Lastly, a gene outside the L locus, *neurobeachin* or *rugose,* which is implicated in altitude-based differences in wing shape between *Heliconius* species^48,49^, and also needed for the formation of short-term olfactory aversive memory in *Drosophila*^50,51^, is downregulated in *H. m. malleti* brain compared to *H. m. rosina* brain in the trained conditions (HMEL010907g1, log2FC = −0.13).

## Discussion

Here, we identified transcriptomic differences between sexually mature males of two allopatric *Heliconius melpomene* subspecies (*H. m. malleti* and *H. m. rosina*) that differ in aversive courtship learning^18^. We found that *H. m. malleti* and *H. m. rosina* males are inherently different in their neurotranscriptomic signatures across all three tissues, but had the greatest neurotranscriptomic changes in the brain after a social exposure with a female, suggesting a large role of CNS processing in driving variance in response to a social learning event. Several learning and memory genes were differentially expressed in the brain during training, supporting the heightened involvement of these genes during the mate preference learning process. We report genes/loci that control “magic” traits (wing color, pattern, shape)^21,23,24,26,30,33,37,48,49^, sexual traits (pheromones)^28,45–47^, and preferences^26,30,31,33^ as DE between the two subspecies, and these genes may be simultaneously influencing both preference and preferred traits in a species famous for its assortative mating^26,30,31^. A number of these genes were also differentially expressed between the brains of *H. melpomene* and *H. cydno* during mate choice^31^, suggesting that they are important for both within and between species mate recognition. Together, this provides evidence for the role of putative magic loci, and pleiotropic genes which may influence mate choice, and drive speciation via sexual selection^30,31,33,36,37,52^.

We found transcriptomic differences between the males of the two subspecies in all three tissues, possibly due to inherent differences in geography, ecological conditions, and evolutionary pressures^53^. Visual centers of the brain have diverged in size in sympatric *H. cydno* and *H. melpomene* that occupy different microhabitats^54^. Our observed gene expression differences suggest divergence between these two allopatric subspecies, potentially driven by habitat-specific natural selection. But, we found greater DEGs in the brain, followed by the eyes and the antennae, during a social exposure with a female. This suggests greater CNS-level differences in neural processing between individuals that differ in preference learning ability, similar to that of female paper wasps *Polistes fuscatus*^55^, but different from guppies^56^ and the butterfly *Bicyclus anynana*^34,57^. However, our study on *H. melpomene* differs from the studies involving wasps, guppies, and *B. anynana*, as we were testing the genetic mechanisms responsible for the differences in mate preference learning ability (where both good and bad learners are given a learning opportunity), and not preference learning or its valence itself (which includes naive and trained individuals). Higher processing in the brain may be responsible for the variation in preference learning abilities between individuals, whereas sensory processing may be more important when individuals are learning different traits/trait values.

Though the brain had greater DEGs associated with differences in mate preference learning, we identified many known learning genes and learning associated GO terms in all three tissues in our study. These include *neurobeachin,* which is required in mushroom body development, olfactory learning, and short-term memory consolidation^50,51^, but is also implicated in altitude-based wing shape variations between *Heliconius* species^48,49^, and *vitellogenin*, which is implicated in social behaviors and is DE in the butterfly *Bicyclus anynana* during a social imprinting-like learning^34,58–60^. We found gene-network modules enriched for zinc ion binding to be associated with variance in preference learning in all three tissues between the two subspecies. Zinc plays a role in synaptic plasticity and signaling, activating signaling cascades that are required for learning and memory formation^39,40^. Dietary deficiency in zinc is known to reduce memory retention in mice^61,62^. Thus, many genes identified in our study are implicated in learning across species, suggesting a conserved, context-independent learning pathway.

*H. melpomene* subspecies differ in a few genomic “hotspots” that are under natural and sexual selection^19–21^. These “hotspots” regulate “magic” traits such as wing color, pattern, and shape, sexual traits such as pheromones, and their preferences. Many genes adjacent to wing patterning genes were DE between the two subspecies in various scenarios. These include the visual mate preference genes *regucalcin 1*, *regucalcin 2*, and *GRIK2* within the *optix* wing patterning gene locus^26,30,31^. *Regucalcin 1* perturbation disrupts *Heliconius* males’ ability to court females^26^. *Regucalcin* regulates *Ca^+2^/Calmodulin-dependent protein kinase II (CaMKII)* gene activity, which is implicated in learning and memory acquisition^63^ whereas, *GRIK2* binds to molecules like NMDA (N-methyl d-aspartate), that regulates olfactory learning in *Drosophila*^64^, social behavior and female mate preferences in swordtails, *Xiphophorus spp.*^65,66^, and guppies, *Poecilia reticulata*^56,67^. *Wash* (control) and *dome* (control and trained)–within the *cortex* wing patterning locus–are DE between the two subspecies in the brain. *Wash* is necessary during oogenesis^68^ while *dome* is implicated in long-term olfactory aversive learning and memory in *Drosophila melanoaster*^69^. Lastly, we identified DEGs which are also associated with locomotion variation in these two subspecies^21,48,49^. Two genes (*Col4α1, Vkg*) implicated in the differences in wing beat frequencies between *H. m. malleti* and *H. m. rosina*^21^ are also upregulated in trained *H. m. malleti* brains in our study. This suggests that allelic variation at *Col4α1* and *Vkg* could simultaneously influence flight and mate selection behavior.

*Heliconius* also exhibit diverse pheromones^28,45^ and we found two DEGs (*Alcohol dehydrogenase* and *fatty acyl-CoA reductase*) within the putative QTL region of chromosome 20 associated with male pheromone production in *H. melpomene*^47^. These genes are associated with differences in male androconial pheromone production between *H. melpomene* and *H. cydno*^47^. Other genes include *GGPS1* which acts as an anti-aphrodisiac that males transfer to the females during mating, to reduce female remating^46^. All three genes were DE between the two subspecies during training, suggesting that these genes may control both the pheromone production and preference for odor cues. Future studies could decipher the association of these genes to courtship and mating related odor preferences, similar to what has been achieved for visual preference genes^26,30,31^.

The retrieval of many DEGs within putative “magic” loci across sensory modalities suggests that assortative mating in *Heliconius* butterflies may be facilitated by a suite of genes and loci with pleiotropic effects on preference and preferred traits (Figure 5). Selection on these loci can drive the evolution of mate preference learning due to linkage, and social conditions like mate rejection can activate genes on such loci leading to an aversive learning experience (Figure 5). Yet, in lineages like *H. m. rosina*, mate rejection may repress learning genes as observed in this study, thereby producing a lack of learned change in behaviors (Figure 5). Many of these genes are also implicated in neural signaling, learning, and memory formation. Future studies should investigate the dynamics of such pleiotropic loci across different lineages of *Heliconius* to understand the evolutionary patterns of mate preference learning.

**Figure 5:**
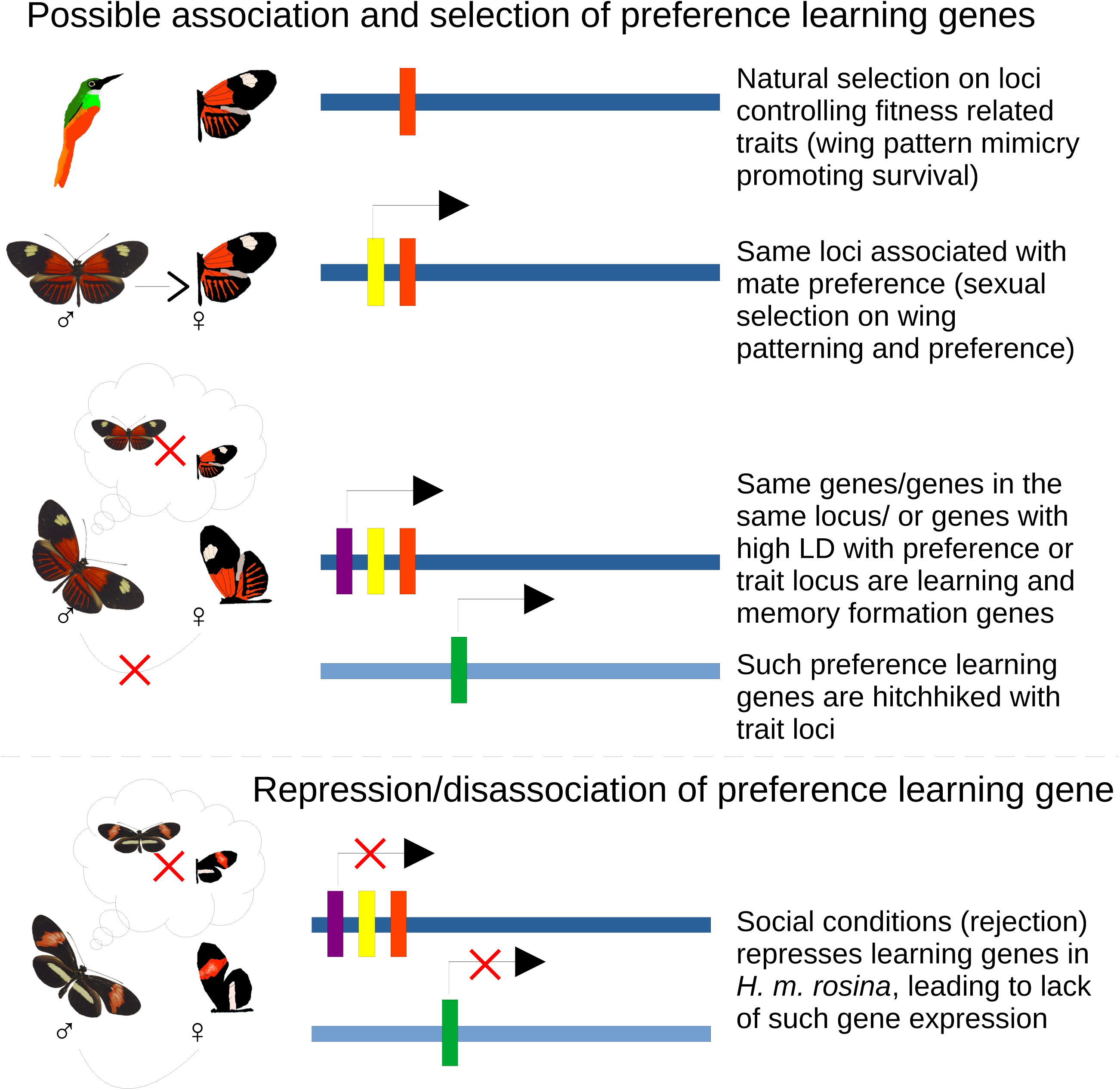
Mate preference learning genes may be pleiotropic, or tightly linked to naturally (or sexually) selected trait genes: (Top to bottom) Genes regulating naturally (or sexually) selected traits (such as wing patterning; orange vertical box) may be adjacent or in high linkage disequilibrium (LD) with genes associated with mate preferences (yellow vertical box). These same genes/loci could have pleiotropic functions for learning and memory formation, or they could be adjacent to the sexually selected trait/preference genes (violet vertical box), or they could be in a different location, but in high LD with the trait or the preference locus (green vertical box). These learning genes, either present adjacent to, or in high LD with positively selected (due to natural or sexual selection) trait/preference loci, could get hitchhiked during recombination. Social learning condition, such as mate rejection, could activate such learning and memory formation genes, subsequently affecting the observed reduced courtships in *H. m. malleti* males^18^. On the contrary, social conditions (mate rejection) may repress learning and memory genes, thereby inhibiting gene expression and downstream lack of mate preference learning in *H. m. rosina* males^18^. In this figure, horizontal longer bars (dark and light blue) are chromosomes, and vertical shorter bars are representative genes. Arrows indicate activation of genes leading to their expression, and red cross (X) on arrows indicate repression of genes leading to lack of their expression. In the cartoons, the bird is Rufous-tailed jacamar, and the butterflies above dashed line are *H. m. malleti* and butterflies below the dashed line are *H. m. rosina*. (All depictions are not to scale).

## Conclusions

In this study, we set out to identify the neurotranscriptomic mechanisms underlying natural differences in male mate preference learning between two allopatric populations of *Heliconius* butterflies. While there were consistent neurotranscriptomic differences between the two subspecies across all three tissues, we identified greater gene expression in the brain compared to the sensory tissues during preference learning, suggesting a heightened role for CNS to incorporate learned preferences. Lastly, we identified DEGs within putative “magic” loci that control both the development of sexual traits, preference for wing color patterning, and locomotory behaviors, as well as genes that control pheromone signals. Most of these putative “magic” genes are also involved in neural development, learning, and memory acquisition. Taken together, these pleiotropic genes could be responsible for three-way functions across spatial and temporal scales in an individual, and selection (natural and/or sexual) on such genes in the population may drive divergence in behaviors such as mate preference learning.

## Materials and Methods

### Study species and husbandry

*Heliconius melpomene* (Family: Nymphalidae) is a Central and South American butterfly with a wide distributional range, from Nicaragua in the north to southern Brazil in the south^53^. Live pupae of these two phenotypes were shipped from Ecodecision Heliconius Works in Ecuador, and Costa Rica Entomological Supply in Costa Rica to the butterfly greenhouse facility in University of Arkansas, which is maintained at ∼27°C, average relative humidity of 70% and a 13:11 hour L:D cycle. Once the butterflies emerged from their pupae, they were tagged using a silver sharpie (SHARPIE 39108PP), and sexed. Males were housed in sex- and phenotype-specific cages (60.96 x 60.96 x 142.24 cm), while females of both *H. m. malleti* and *H. m. rosina* were housed together. Both the male and female cages were visually isolated from each other, with males and females having no exposure to the opposite sex till the day of the behavioral assay. All butterflies were fed *ad libitum* with BIRDS choice butterfly nectar (Birdschoice, Chilton, WI, USA) and pollen from *Lantana spp* flowers.

### Behavioral assays for head collections

Behavioral assays were conducted from 11:00 to 12:30. To replicate the experience that induces a change in behavior in *H. m. malleti* but not in *H. m. rosina*^18^ (Figure 2A), hereafter called the training treatment, a 10-day-old naïve male was exposed to a 3-5-day old female of the same phenotype for 90 minutes in an observation cage (60.96 x 60.96 x 142.24 cm). The males were allowed to court the females in those 90 minutes, but not allowed to mate with the females. If the males courted, we recorded that and removed the female from the observation cage before copulation. During the control treatment, in each assay, a 10-day-old naïve male was housed alone in the observation cage for 90 minutes, without any female exposure. Cages were separated by white particle board, so individuals could not see into other cages and/or treatments. After 90 minutes in either the training or the control treatment, the males’ heads were decapitated using an RNA-free dissection scissors, and immediately flash frozen in liquid nitrogen and stored at −80°C till RNA extraction. We collected 10 samples for trained *H. m. malleti* treatment, 9 samples for trained *H. m. rosina* treatment, 11 samples for control *H. m. malleti* treatment, and 11 samples for control *H. m. rosina* treatment (Figure S1). Heads were collected from March 2019-November 2020.

### RNA extraction, cDNA library preparation and sequencing

We dissected the heads in RNA-later ICE (Ambion; Austin, TX, USA) after acclimating and incubating the heads in RNA-later ICE at −20°C for approximately 16 hours. During dissection, we removed and discarded the cuticle, proboscis, and other fatty tissues. We separated the antennae (AN), eyes (EY), and the brain (BR) from other tissues with as little contamination as possible and disrupted them mechanically with a pestle in lysis buffer. We extracted large RNA (>200 nucleotides) for the three tissues separately using the Nucleospin® miRNA kit (Macherey-Nagel; Düren, Germany) following the manufacturer’s protocol, and assessed RNA quantity and quality using NanoDrop 2000 (Thermo Fisher Scientific; Waltham, MA, USA) and Tapestation 2200 (Agilent; Santa Clara, CA, USA) before preparing cDNA libraries.

We prepared the cDNA libraries with a 200ng starting RNA concentration using KAPA mRNA HyperPrep Kit and Unique Dual-Indexed Adapters (KAPA Biosystems; Wilmington, MA, USA) following the manufacturer’s protocol. Library quality was analyzed by NanoDrop 2000 and Tapestation 2200, followed by subsequent assessment on a 5300 Fragment Analyzer (Agilent; Santa Clara, CA, USA). Libraries were sent to the University of Chicago Genomics Center, where they were sequenced on an Illumina NovaSeq 6000, with 100bp single-end sequencing. The AN libraries were sequenced separately whereas the EY and BR libraries were sequenced together on a single flowcell.

#### Reads quality control, adapter trimming, and alignment

The quality of all the raw reads obtained from the University of Chicago Genomics were checked using FastQC v0.11.5 (http://www.bioinformatics.babraham.ac.uk/projects/fastqc/). All samples passed the initial QC check. We used Trimmomatic v0.38 to trim the Illumina adapter sequences from the raw reads^70^. We used STAR v2.7.4a to create a *de novo* genome index for *H. melpomene* from the *H. melpomene melpomene* v 2.5 genome downloaded from Lepbase (http://ensembl.lepbase.org/Heliconius_melpomene_melpomene_hmel25/Info/Index)^71^. We aligned the adapter-trimmed reads to the *de novo* genome index using STAR v2.7.4a using the “--twopassMode Basic” option. The percentage of sequences aligned to the genome ranged from 24.11% to 92.06% and varied between tissues, with the best alignment for EY tissues, followed by BR and AN (Figure S2A; Table S1). Four AN tissues (SP-12, SP-35, SP-36, SP-38) were below 50% unique alignment and were discarded from downstream analyses. The tissues segregate separately in PCA space (Figure S2B).

#### Differential gene expression analyses

We performed differential gene expression analyses for the three tissues (AN, EY, BR) separately using DESeq2 v 1.24.0 package^72^ for R version 4.3.0 (R Foundation for Statistical Computing, Vienna, Austria) ^73^ using a generalized linear model design:

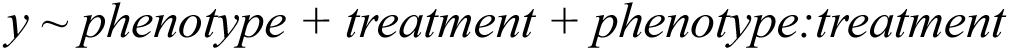

where *y* is gene expression, phenotype is *H. m. malleti* vs *H. m. rosina*, and treatment is control vs training. With this design, we compared gene expression between five different combinations: 1. *malleti* control vs *rosina* control; 2. *malleti* trained vs *rosina* trained; 3. *malleti* control vs *malleti* trained; 4. *rosina* control vs *rosina* trained; and 5. the interaction between phenotype and treatment. We only used genes that had ≥10 total mapped reads, and a gene was considered to be DE if its log2FC passed the false discovery rate (FDR) of <0.05 using the “apeglm” package in R^74,75^. We created heatmaps for the above combinations using “pheatmap” v1.0.12 package in R^76^.

#### Functional annotation and GO enrichment analyses

We downloaded the *Heliconius melpomene melpomene Hmel.2.5*^77^ version of the genome from Lepbase (lepbase.org) and extracted the gene IDs and corresponding gene sequences using getfasta script from bedtools v2.31.0^78^. The “genes-only” fasta file was then divided into 1000 gene sequences per fasta file using the perl script of fasta-splitter-0.2.6^79^. We used these 1000 gene-divided fasta files as query and identified the top 10 blast hits blasted against the NCBI “nr” database using the blastx function from DIAMOND 2.1.7 with an evalue of 1e-3 and saved them in a xml file^80^. We performed the annotation and mapping steps using these xml best blast hit files using Blast2GO 6.0.3 software using default settings^81^. We performed GO enrichment analysis using Fisher’s exact test by using FDR<0.05 for multiple testing in the “FilterMode”. We used all the expressed genes as the reference set and DEGs obtained from DESeq2 as the test set. We also performed GO enrichment analyses on co-expression modules obtained from WGCNA analysis by using genes from a particular module as the test set and all the genes expressed as the reference set.

#### Weighted Gene Co-expression Network Analyses (WGCNA)

We performed WGCNA to identify correlated gene networks based on their expression data for six different treatment combinations: malletitrained vs rosinatrained; malleticontrol vs rosinacontrol; malletitrained vs malleticontrol; rosinatrained vs rosinacontrol; malleti vs rosina; trained vs control. We performed these analyses for the three different tissues (BR, EY, AN) separately using the WGCNA version 1.70-3 R package^82^ using similar parameters from^34^. Briefly, we first filtered genes with <10 reads in >90% of the samples and performed “varianceStabilizingTransformation” after running DEseq2. The signed co-expression genes were assembled by setting soft-thresholding power to 7 for brain, 6 for eyes, and 9 for antennae. We then extracted co-expressed modules using the “cutree-Dynamic” function and merged similar modules together using “mergeCloseModules” with “cutHeight” set to 0.25. To obtain module-trait association, we used “binarizeCategoricalVariable” function to create pairwise comparisons of the six different treatment combinations (see above). We adjusted the p-values for each association using the FDR correction method^74^ and any module-trait association with FDR<0.05 were considered significant. We then performed GO enrichment analyses for genes in all the significant modules in the three tissues separately using Blast2GO (see GO enrichment analyses section).

#### Identification of wing patterning, pheromone production, locomotory behaviors, and male mate preference genes

Using our DEGs between *H. m. malleti* and *H. m. rosina,* we set out to explore whether some of the DEGs are known wing patterning genes^83^, pheromone production genes^47^, locomotory behavior genes^21,49^, or male mate preference genes^26,31^. We matched and identified gene descriptions, and sequence IDs described and characterized from these studies to the *H. melpomene* transcriptome, and DEG lists generated in this study, using R statistical software. Using the same method above, we further compared whether DEGs in the brains between *H. m. rosina* and *H. cydno chioneus* during mate preference^31^ were found to be DE in our study between *H. m. malleti* and *H. m. rosina* brains in the control treatment, as we wanted to highlight whether similar genes are involved in mate preference between species of *Heliconius,* and between *H. melpomene* subspecies. Next, we also compared whether DEGs in the developing wings between *H. m. rosina* and *H. m. melpomene*^37^ were found to be DE in our study between *H. m. malleti* and *H. m. rosina* in the control treatment in all three tissues (BR, EY, AN), as these genes would be pleiotropic for wing development and mate preference. For both these analyses, we incorporated the DESeq2 pipeline as described above using RNA sequencing raw reads data from^31,37^.

## Supporting information

Supplementary Results

## Acknowledgments

This research was funded by a Grants-in-Aid of Research (GIAR) award from the Society for Integrative and Comparative Biology (SICB) to SP, an NSF IOS-1937201 to ELW, and the University of Arkansas, Fayetteville, AR USA. We thank the members of the Westerman Lab in butterfly husbandry, and David A. Ernst, and Yi Ting Ter for their help in bioinformatics. We thank Brian A. Counterman, Jeffrey Lewis, William J. Etges, Mehrnaz Afkhami, Michael Sheehan, Colby Behrens, and Ben Sandkam for thoughtful comments and feedback on the manuscript. This research is also supported by the Arkansas High Performance Computing Center which is funded through multiple National Science Foundation grants and the Arkansas Economic Development Commission. We thank Jeff F. Pummill for assistance in using the Core computing facility at the University of Arkansas.

## Source data

Raw reads can be found on NCBI Sequence Read Archive (SRA) database under the BioProject PRJNA1424375. All scripts and a README file to replicate RNAseq data analyses are deposited in a Github repository, and can be accessed using the link: https://github.com/sdpotdar/Heliconius-melpomene-sensory-and-neural-transcriptomics

## Supplementary information

1. Supplementary results: Descriptions of supplementary results (supplementary_results.docx)
2. Supplementary tables: A workbook containing supplementary tables (S1-S41). The first sheet includes table of contents, and the descriptions of each table (supplementary_table.xlsx).

## Extended data figures

**Figure S1:**
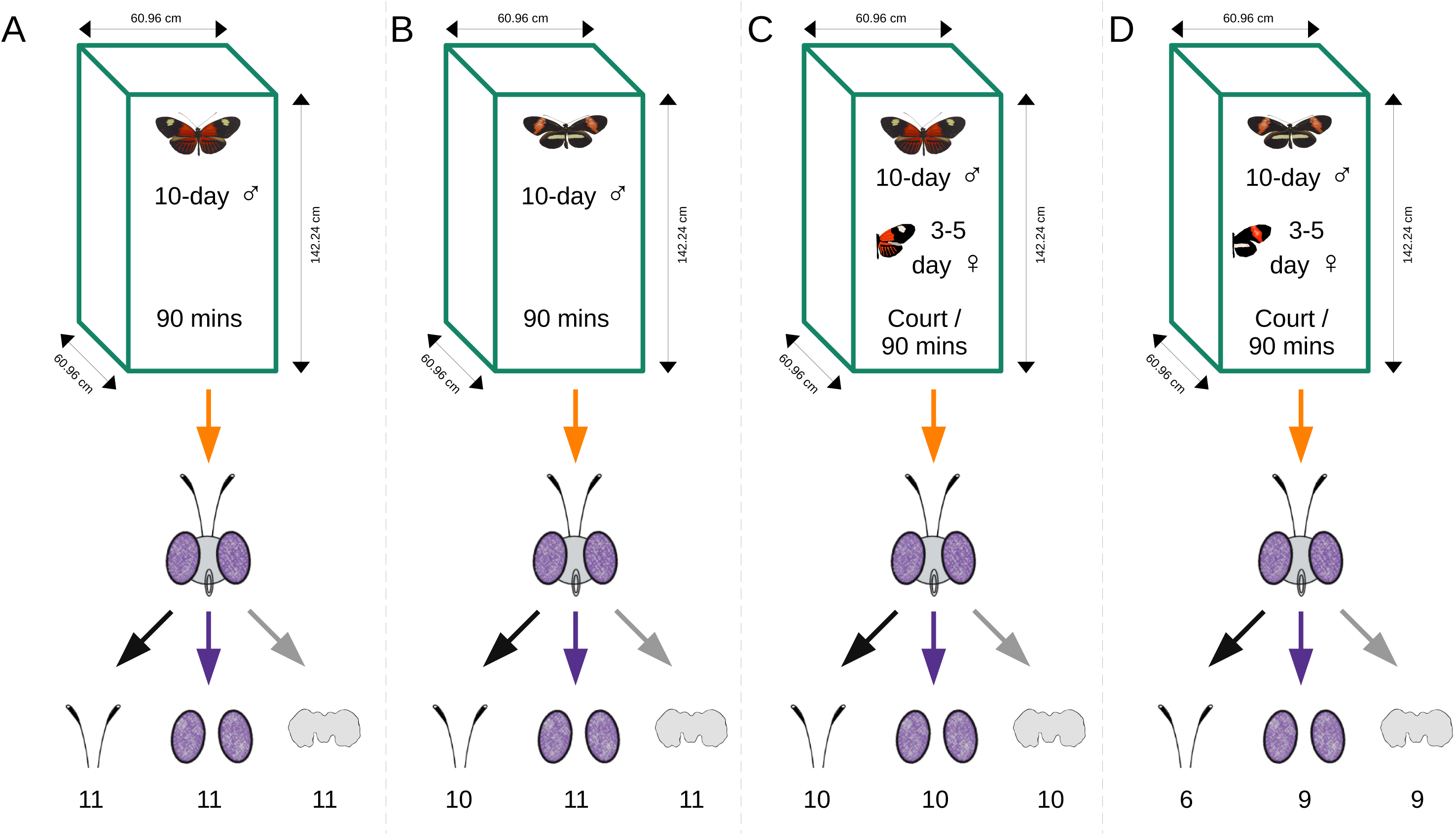
Experimental design to test transcriptomic differences between control, and trained *H. m. malleti* and *H. m. rosina*: 10-day old males of A) *H. m. malleti,* and B) *H. m. rosina* were kept in an experimental cage for 90 minutes, after which the heads were collected and transported in liquid nitrogen, followed by long term storage in −80°C. Antennae (AN), eyes (EY), and the brain (BR) were dissected for RNA extraction, cDNA library preparation, and downstream DESeq analyses. In training treatment, 10-day old males of C) *H. m. malleti* and D) *H. m. rosina* were exposed to 3-5-day old females of their own subspecies, either till the males courted, or for 90 minutes, whichever came first. If the males courted, females were removed before copulation, and the males remained in the experimental cage for 90 minutes. Heads were collected, transported, and stored in −80°C similar to A and B (control conditions). RNA was extracted from AN, EY, and BR for cDNA library preparation, followed by DESeq analyses. The number of tissues dissected for RNA extraction in every treatment is indicated below the tissue symbol.

**Figure S2:**
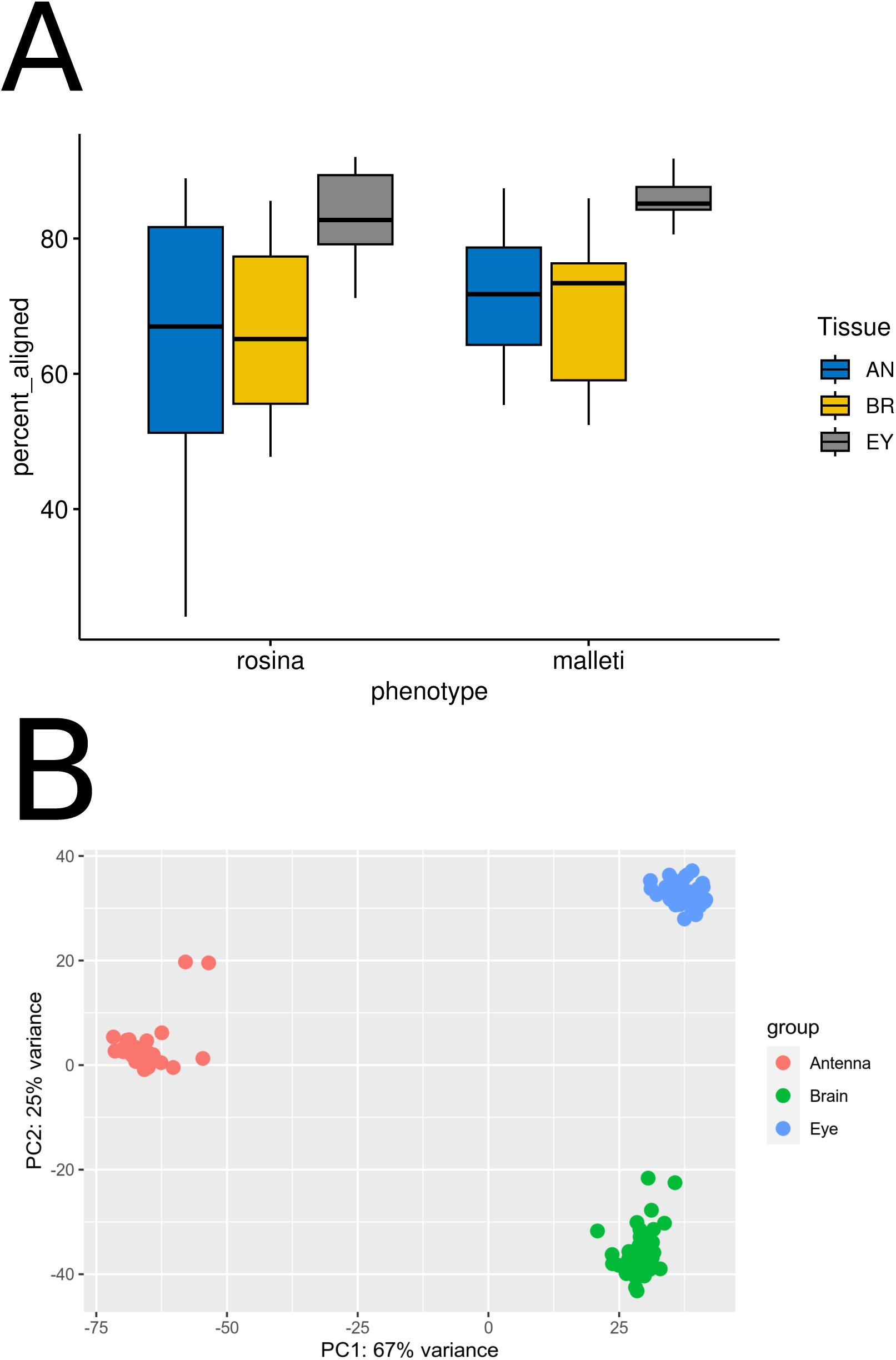
Gene expression segregates by tissues: A) The percentage of sequences from the eyes (EY) tissue had a better alignment to the Hmel 2.5 genome for both *H. m. malleti* and *H. m. rosina*, followed by the brain (BR), and the antennae (AN). Nevertheless, B) gene expression from the three tissues segregated separately in a principal components analysis (PCA) plot, either suggesting different genes are involved, or a stark tissue specific expression patterns of shared genes, or both.

**Figure S3:**
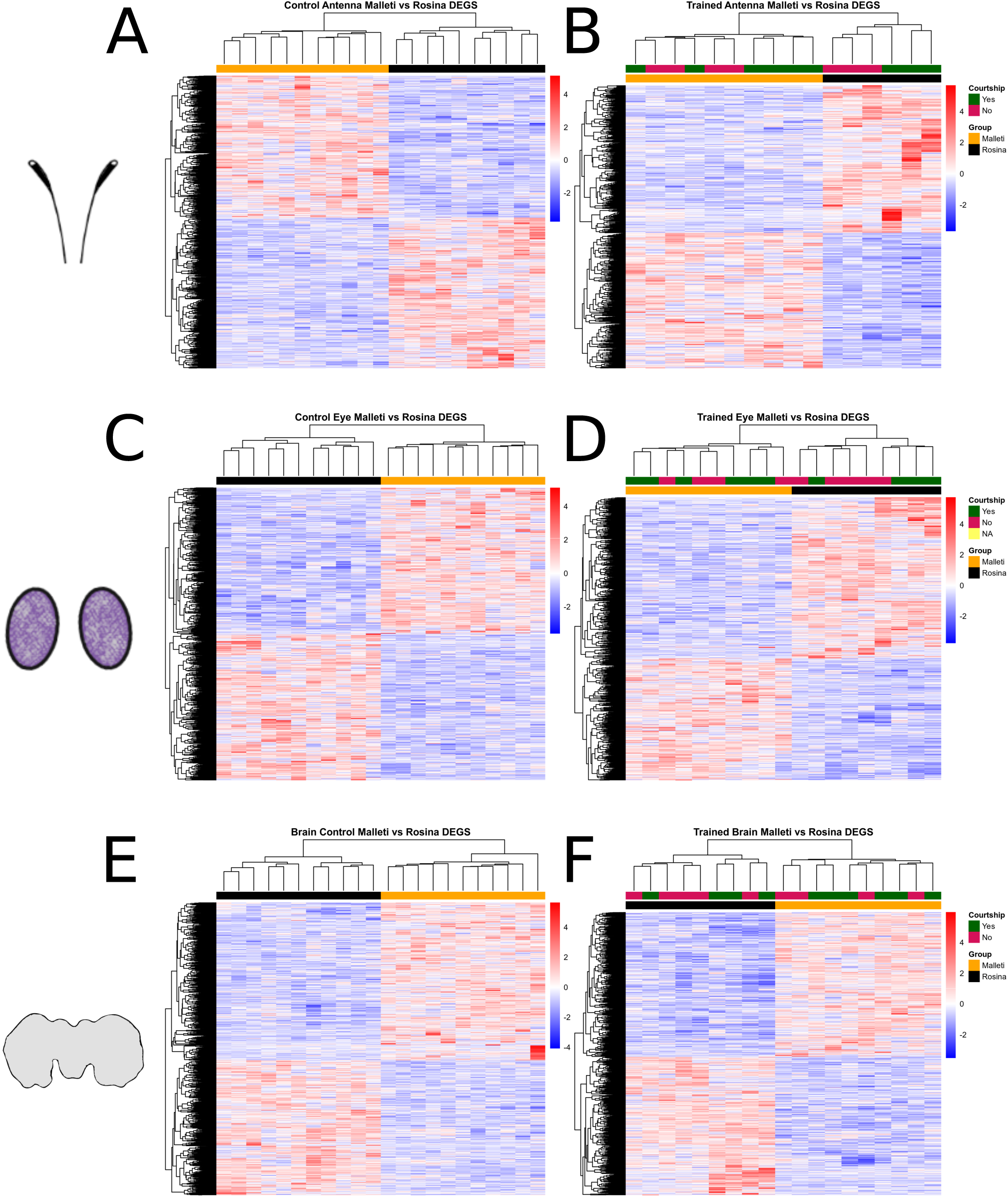
All DEGs in both the control and the training conditions segregate *H. m. malleti* and *H. m. rosina* separately in all three tissues: Heatmaps of all DEGs in A) *H. m. malleti* vs *H. m. rosina* control condition in the antennae; B) *H. m. malleti* vs *H. m. rosina* training condition in the antennae; C) *H. m. malleti* vs *H. m. rosina* control condition in the eyes; D) *H. m. malleti* vs *H. m. rosina* training condition in the eyes; E) *H. m. malleti* vs *H. m. rosina* control condition in the brain; F) *H. m. malleti* vs *H. m. rosina* training condition in the brain. In these heat maps, each row is a single gene, and each column is an individual sample (indicated by cluster trees in both rows and columns). Gene expressions are plotted as Z-scores after variance-stabilizing transformation, with warmer (red) colors indicating increased expression, and cooler (blue) colors indicating decreased expression relative to the mean for that gene. Heatmaps are clustered based on subspecies (*H. m. malleti* and *H. m. rosina*) and sub-clustered by courtship in B, D, and F.

**Figure S4:**
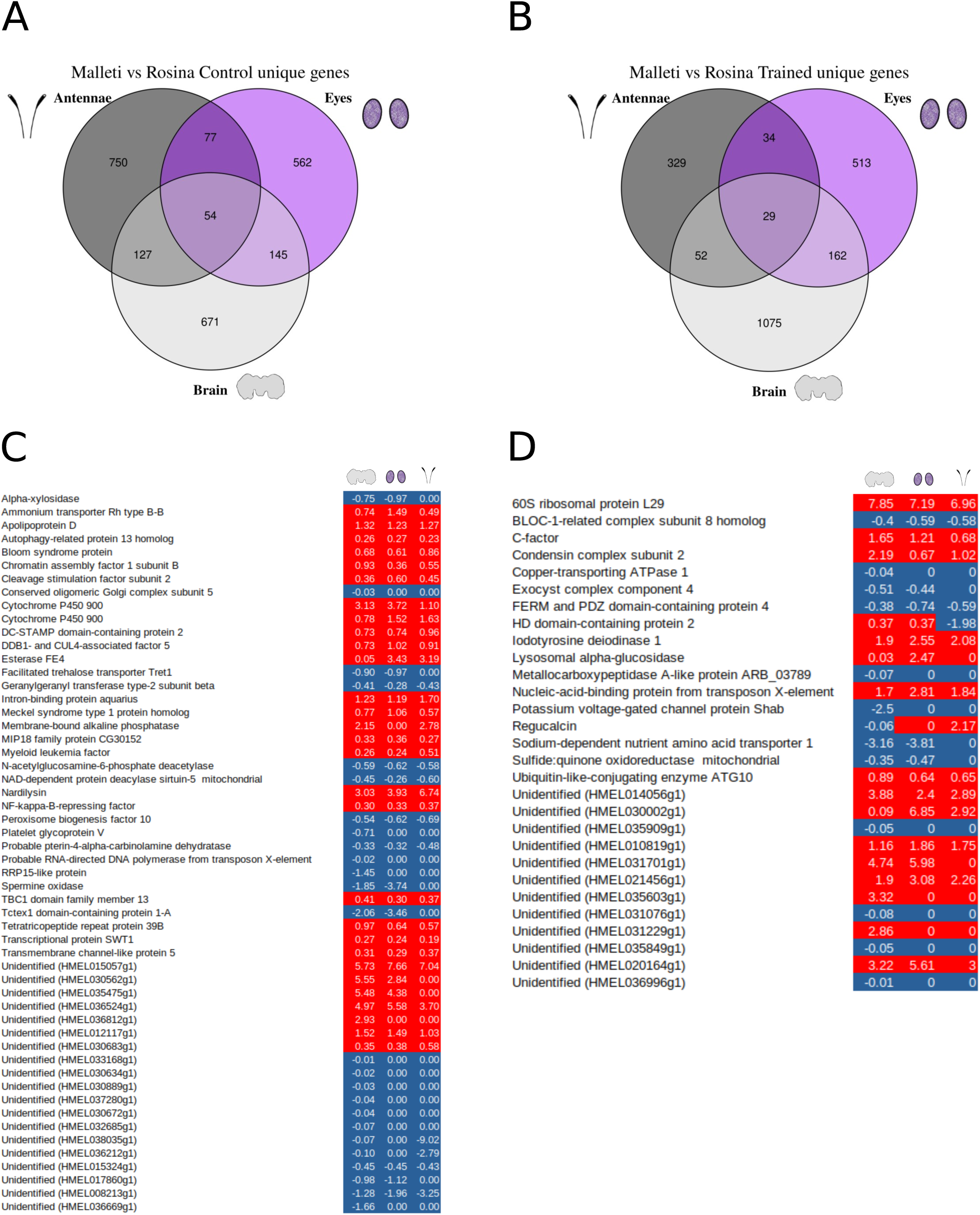
Uniquely treatment specific genes do not change expression patterns across tissues: Unique genes common across the three tissues between *H. m. malleti* and *H. m. rosina* were in A) 54 in the control conditions, and B) 29 in the training conditions. These common genes do not change in their expression patterns across the three tissues between *H. m. malleti* and *H. m. rosina* in both C) the control, and D) the training conditions. In the venn diagrams in A, and B, the numbers indicate unique DEGs for those conditions, in/between those tissues. In C, and D, each row is a gene, whereas each column is the expression of that gene corresponding to the tissues. Red color indicate higher expression, whereas blue color indicates lower expression. Numbers within each cell represents the Z-score. Some cells that have ‘0’ have very low expression that is rounded off, but the colors correspond to their expression patterns (see Table S15 and S16).

**Figure S5:**
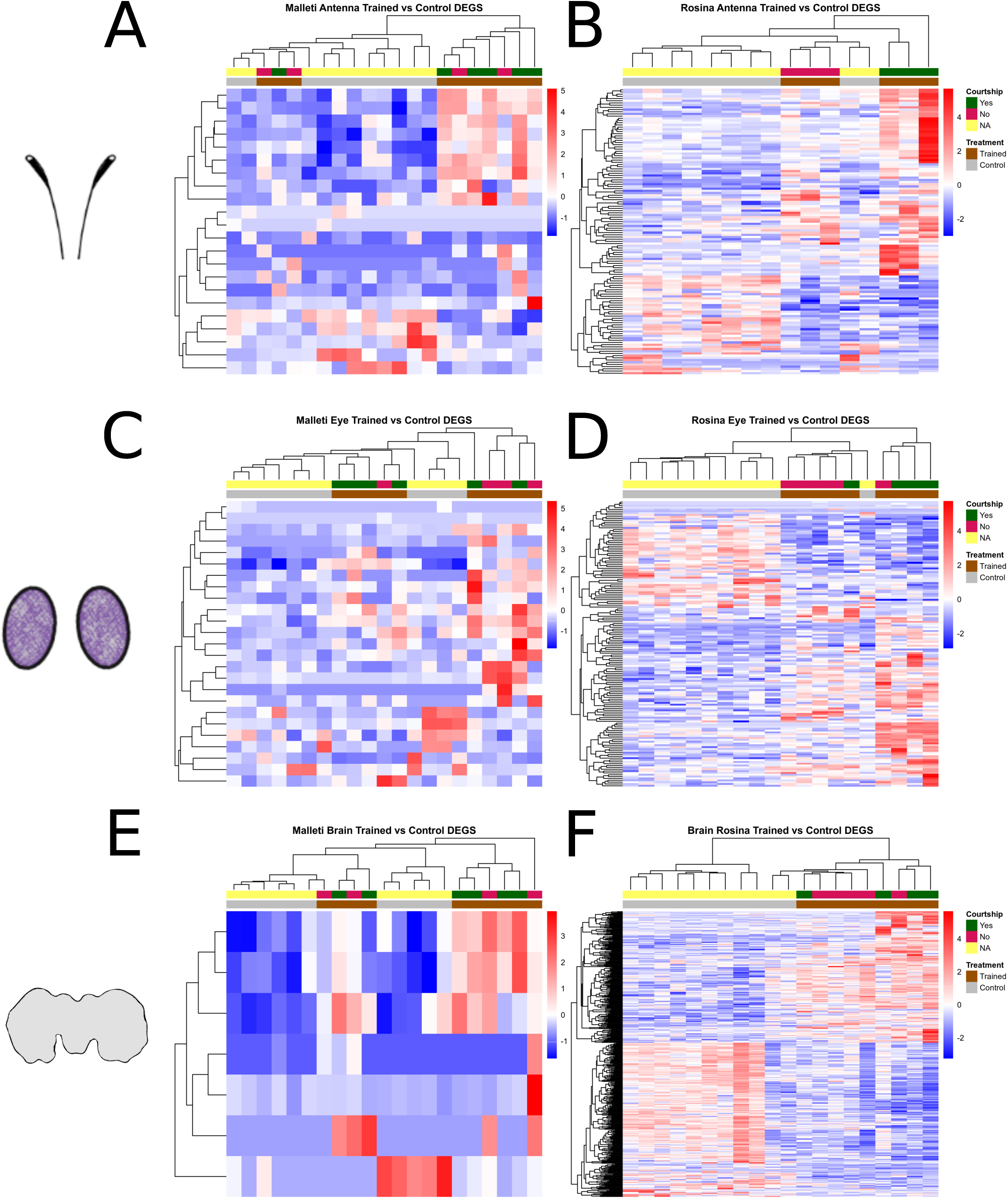
Lesser number of DEGs associated with female exposure in male *H. m. malleti*: When compared between trained and control individuals within subspecies, there were lesser number of DEGs associated with a female exposure in *H. m. malleti* males, compared to *H. m. rosina* males in all three tissues. Heat maps of all DEGs in A) trained vs control condition in *H. m. malleti* antennae; B) trained vs control condition in *H. m. rosina* antennae; C) trained vs control condition in *H. m. malleti* eyes; D) trained vs control condition in *H. m. rosina* eyes; E) trained vs control condition in *H. m. malleti* brain; F) trained vs control condition in *H. m. rosina* brain. In these heat maps, each row is a single gene, and each column is an individual sample (indicated by cluster trees in both rows and columns). Gene expressions are plotted as Z-scores after variance-stabilizing transformation, with warmer (red) colors indicating increased expression, and cooler (blue) colors indicating decreased expression relative to the mean for that gene. Heatmaps are clustered based on subspecies (*H. m. malleti* and *H. m. rosina*) and sub-clustered by courtship data. Note that control individuals do not have courtship data (NA) as they were not exposed to a female.

**Figure S6:**
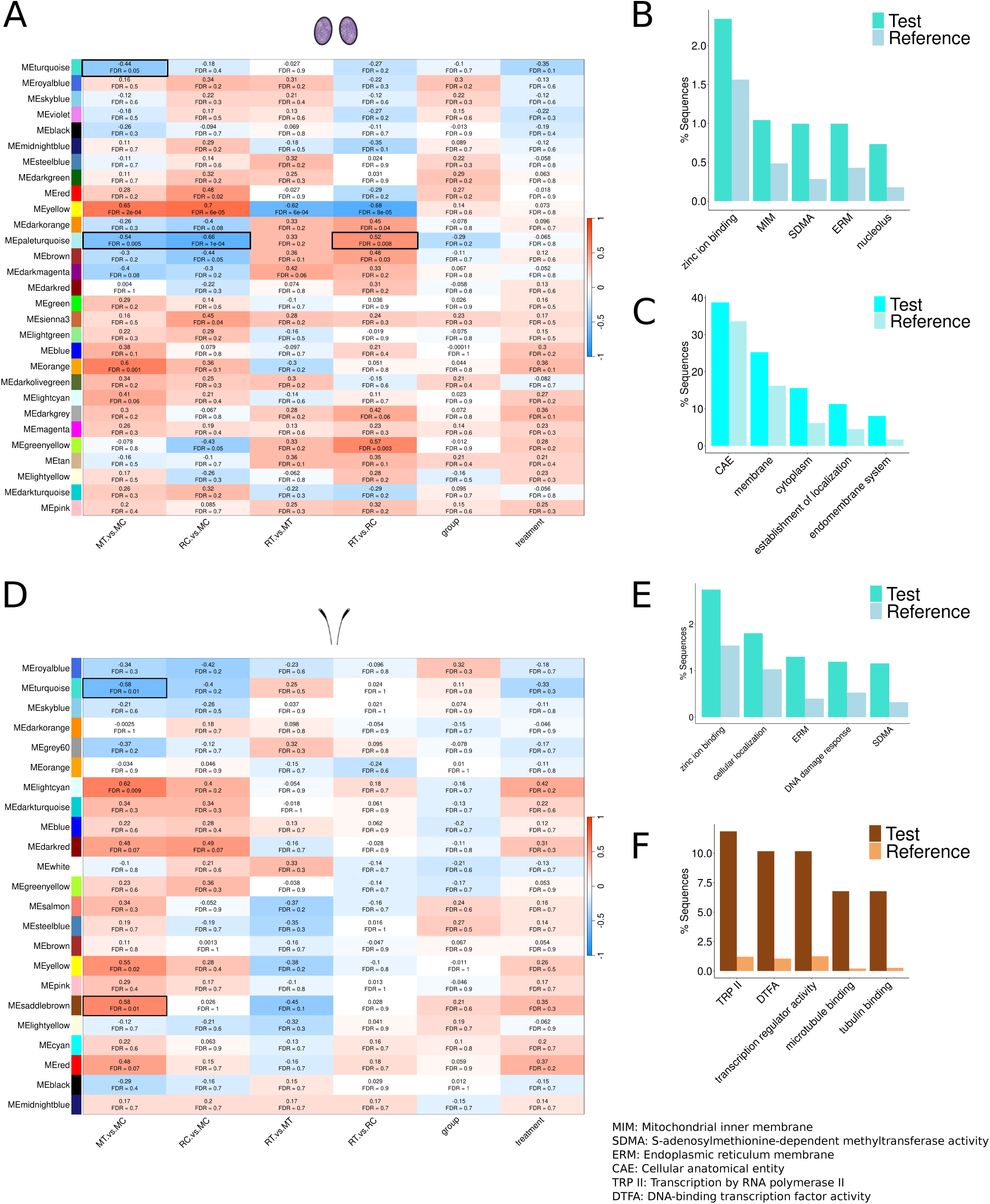
Eye and antennae gene network modules are associated with differences in *H. melpomene* male mate preference learning: Heatmap of gene module-trait association indicates A) two modules (turquoise and orange) that are uniquely associated with female-associated differences in *H. m. malleti* (MT vs MC) in the eyes, while the paleturquoise module is associated with differences between the subspecies. The top enriched gene ontology (GO) terms of B) the eye turquoise module, and C) the eye pale turquoise module. D) Four modules (turquoise, lightcyan, yellow, and saddlebrown) are uniquely associated with female-associated differences in *H. m. malleti* (MT vs MC) in the antennae. The top enriched GO terms of E) the antennae turquoise module, and F) the antennae saddlebrown module. In these heatmaps (A, D), each row is a module eigengenes (ME), whereas each column is a trait comparison. In each box, the strength of association (from −1 to 1) and the false discovery rate (FDR) are reported. MT vs MC = *H. m. malleti* trained vs *H. m. malleti* control; RC vs MC = *H. m. rosina* control vs *H. m. malleti* control; RT vs MT = *H. m. rosina* trained vs *H. m. malleti* trained; RT vs RC = *H. m. rosina* trained vs *H. m. rosina* control; group = *H. m. malleti* vs *H. m. rosina*; treatment = trained vs control. In (B, C, E, F), the test set contains percentage of sequences annotated for that term among all the genes in the module, and the reference set contains the percentage of sequences annotated for that term in the transcriptome.

**Figure S7:**
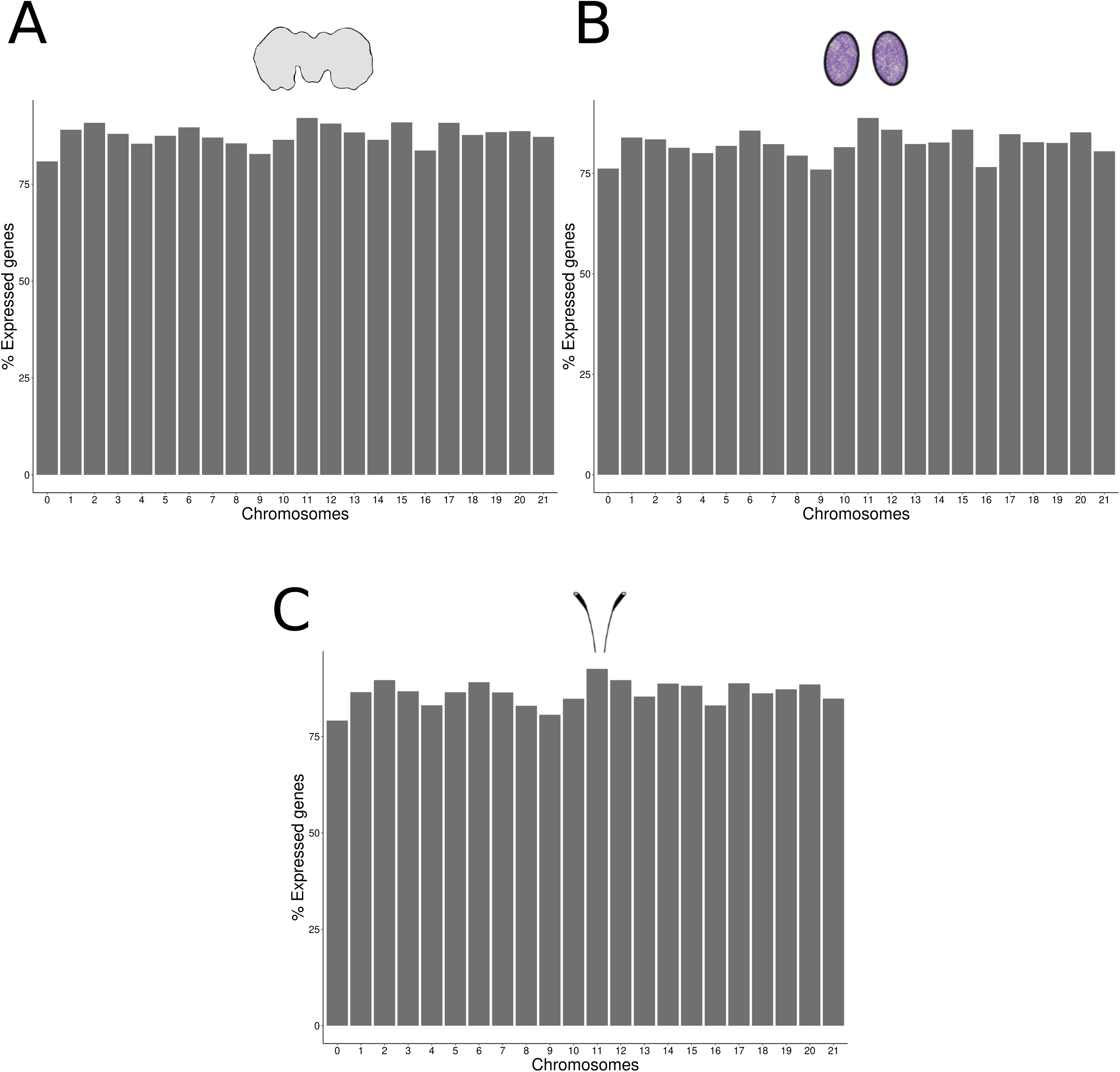
Genes are uniformly expressed across all chromosomes in all three tissues: Barplots depicting the percentage of genes expressed in the A) brain; B) eyes; and the C) antennae are uniformly distributed across all chromosomes in the *Heliconius melpomene* genome. Genes represented on chromosome 0 have not been assigned to any specific chromosome in the genome.

**Figure S8:**
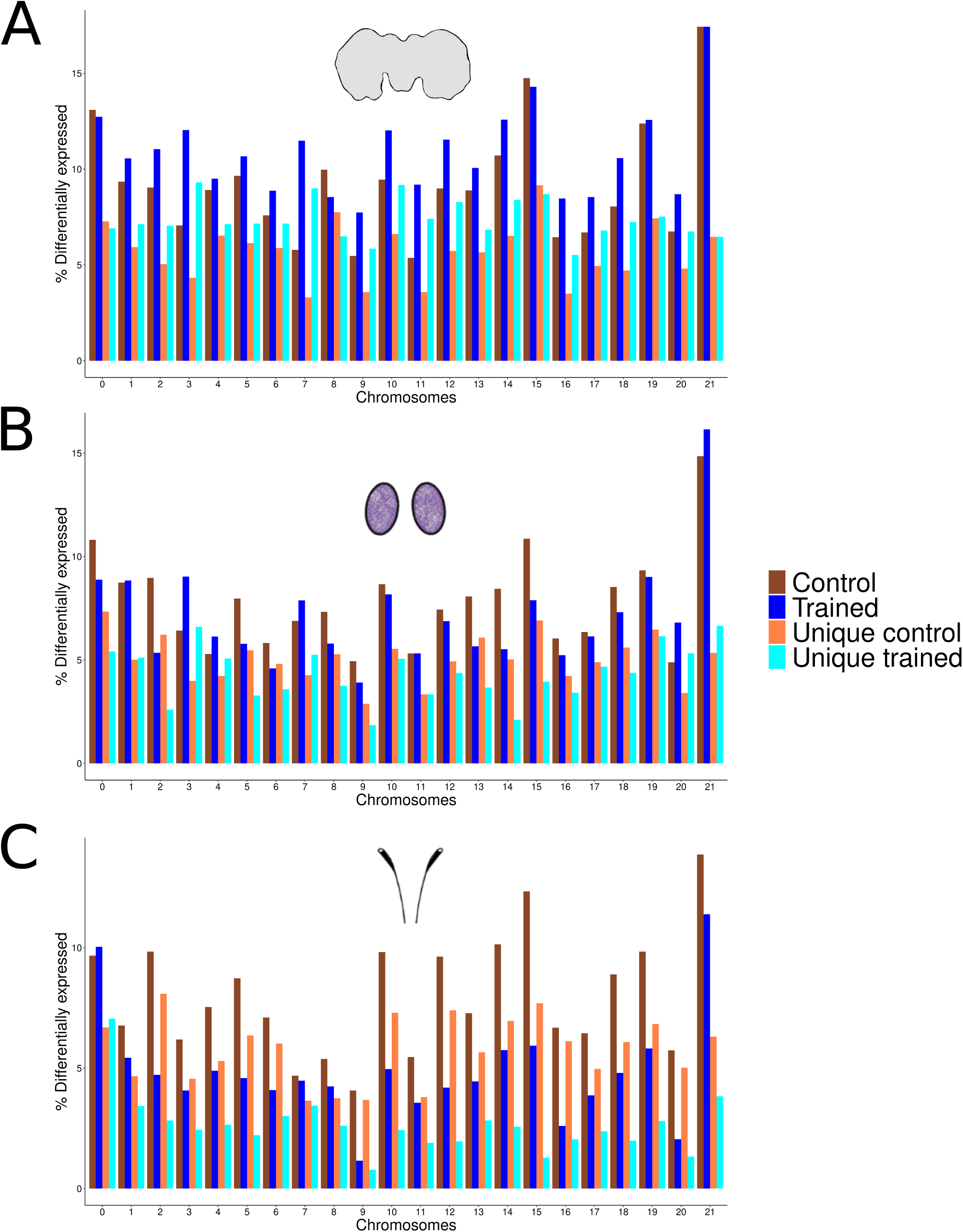
Z chromosome is biased towards DEGs between *H. m. malleti* and *H. m. rosina,* irrespective of social conditions: Percentage of differentially expressed genes between *H. m. malleti* and *H. m. rosina* across chromosomes of the *Heliconius melpomene* genome either in control (dark and light brown) or in trained (dark and light blue) conditions in the A) brain; B) eyes and the C) antennae. Genes represented on chromosome 0 have not been assigned to any specific chromosome in the genome.

