## Supplementary Results for "Neurotranscriptomic signatures of natural variation in mate preference learning in two subspecies of *Heliconius melpomene* butterflies"

***H. m. malleti* and *H. m. rosina* have baseline transcriptomic differences:**

While observed DEGs in free flying isolated males of *H. m. malleti* and *H. m. rosina* are associated with a variety of different biological processes, we were interested in genes associated with mate selection, reproduction, fertilization, and social interactions. Some notable genes associated with reproduction and fertilization and uniquely DEG in the brain include *protein sneaky* (*snky*, HMEL032541g1, log2foldchange = 1.29), *ovochymase-2* (*Ovch-2*, HMEL036161g1, log2foldchange = 1.45), two uncharacterized genes with best blast hit for ejaculatory bulb-specific protein (HMEL003116, log2foldchange = 1.34) and *protein takeout-like* (HMEL014930g1, log2foldchange = 1.46), all of which were upregulated in *H. m. malleti* relative to *H. m. rosina* control brains. These genes are required for fertilization and male courtship behaviors in other insect species^1–4^. While Gene Ontology (GO) enrichment analysis of brain DEGs did not recover any significantly enriched GO terms. Weighted Gene Co-expression Network Analysis (WGCNA) identified two modules (turquoise and greenyellow) associated with baseline differences between the brain transcriptomes of the two subspecies. GO enrichment on the genes in the turquoise module revealed 104 GO terms, some of which were enriched for “neuron differentiation”, “axon guidance”, “signal transduction”, “neurogenesis”, and “response to stimulus” (Table S25).

We also found genes that were implicated in sexual reproduction and fertilization and were upregulated in the eye tissue of control *H. m. malleti* relative to control *H. m. rosina* males. These genes include a *putative vitellogenin receptor* (*yl*, HMEL031407g1, log2foldchange = 1.24), and transcript with best blast hit for a seminal fluid protein (HMEL036651g1, log2foldchange = 4.16); as well as two learning genes (**cAMP-specific 3',5'-cyclic phosphodiesterase, *dnc*, HMEL007752g1, log2foldchange = 0.22; and *henna, Hn,* HMEL011020g1, log2foldchange = 0.46)**. While WGCNA recovered many modules that were shared between control and training conditions (Table S28), only the red module (848 genes) with an unknown hub gene (HMEL011815g1), and the sienna3 module (52 genes) with *aquaporin AQPAE.e-like* (HMEL002342g3) hub gene were uniquely associated with baseline differences between the eyes of *H. m. malleti* and *H. m. rosina* (Figure 6A)*.* GO enrichment analyses for all the DEGs in the eyes, as well as the red and sienna3 modules did not identify any significantly enriched GO terms.

Transcriptional differences in isolated *H. m. malleti* and *H. m. rosina* antennae also included DEGs associated with sexual reproduction and fertilization. Most notable were protein *takeout* (HMEL014783g1, log2foldchange = 2.08), *ovochymase-2* (*Ovch-2*, HMEL036161g1, log2foldchange = 1.92), and an uncharacterized gene which had a best blast hit for a seminal fluid protein (HMEL036651g1, log2foldchange = 1.5), all of which were up-regulated in *H. m. malleti*. GO enrichment of antennae DEGs revealed three significantly enriched GO terms: “oxidoreductase activity”, “carbohydrate metabolic process”, and “helicase activity” (Table S8).

Overall, we found a number of distinct tissue-specific transcriptomic differences between the brains, eyes, and antennae of isolated, free flying *H. m. malleti* and *H. m. rosina* butterflies which may reflect habitat, geographical, and/or other selective pressure differences between these two subspecies. Moreover, identification of two unique WGCNA modules associated with the differences between *H. m. malleti* and *H. m. rosina* in the eyes may suggest divergence in visual centers.

***Variation in learned response is associated with more DEGs in the brain than in sensory tissues*:**

Some of the unique DEGs in the brain include known learning and memory genes and genes that are known to have neural processing or neurodevelopmental functions. For example, *cyclic nucleotide-binding domain-containing protein 2* (*Cnbd2*, HMEL016677g1, log2foldchange = 1.9) and *Tachykinin-like peptides receptor 99D* (*TkR99D*, HMEL031908g1, log2foldchange = 0.41) are upregulated in *H. m. malleti* relative to *H. m. rosina* during the training condition. Apart from learning, memory, and neural signaling genes, we also recovered genes implicated in sexual behavior and male courtship in other systems. The *takeout* gene (HMEL015228g1, log2foldchange = 0.80), which is required for male courtship in *Drosophila melanogaster*^4^, is upregulated in trained *H. m. malleti* brains. A second *takeout*-like ortholog is DE between lineages in the control condition. Other genes that are DE have functions in neurodevelopment, hormone regulation, zinc ion binding, and sensory perception of odor and vision (Figure 2B, C).

In the eyes, we identified 31 DEGs with functions in vision and eye development (Figure 2D), of which 5 were upregulated in *H. m. malleti*. For example, *enabled* (*ena*, HMEL011300g1, log2foldchange=0.36), and *arganine N-methyltransferase 3* (*PRMT3,* HMEL017262g1, log2foldchange = 0.86) were upregulated in *H. m. malleti* during training while two orthologs of *carotenoid isomerooxygenase* (*ninaB*, HMEL034898g1, log2foldchange= -2.14; HMEL034896g1, log2foldchange= -9.4Xe^-6^) were downregulated in *H. m. malleti* during training*.* Learning and memory formation genes including *Cnbd2* (HMEL016677g1, log2foldchange=2.36), *ecdysoneless* (*ecd,* HMEL006646g1, log2foldchange=0.64), and *neuroligin-4, Y-linked* (HMEL012004g1, log2foldchange=0.50) were upregulated in trained *H. m. malleti* eyes relative to *H. m. rosina* eyes.

In the antennae during the training treatment, three known learning genes were DE between *H. m. malleti* and *H. m. rosina* (Figure 2E)*.* These were *henna* (*hn,* HMEL011020g1, log2foldchange = 2.80), *Glutamate NMDA receptor subunit 1*(*Nmdar1*, HMEL017595g1, log2foldchange = 6.7Xe^-7^), and *transient receptor potential protein* (*trp*, HMEL032237g1, log2foldchange = -1.11).

***Training exposure induces large transcriptomic response in H. m. rosina, in spite of missing behavioral response:***

When we compared DEGs between trained and control individuals within each subspecies (*H. m. malleti/H. m. rosina*), we recovered more DEGs in *H. m. rosina* males, who do not exhibit a significant behavioral response to training, than in *H. m. malleti* males, who do (Figure 2B, Figure S5)*.* For *H. m. malleti* we recovered 7 DEGs in brain (Table S18, Figure S5E), 25 DEGs in eyes (Table S19, Figure S5C), and 22 DEGs in antennae (Table S20, Figure S5A). This is in contrast to *H. m. rosina* males, which had 454 DEGs in brain (Table S21, Figure S5F), 155 DEGs in eyes (Table S22, Figure S5D), and 130 DEGs in antennae (Table S23, Figure S5B). Two genes that are implicated in differences in wing beat frequency between *H. m. malleti* and *H. m. rosina* (see main text) are upregulated in trained *H. m. malleti* antennae relative to the antennae of the control individuals. Interestingly, we found many opsins and vision related genes to be upregulated in the antennae of trained *H. m. rosina* individuals relative to the antennae of the control *H. m. rosina* individuals, though this was not the case for *H. m. malleti*. Although we recovered very few DEGs between trained and control *H. m. malleti,* WGCNA identified many modules in all three tissues associated with training in this lineage (Figure 3A, Figure S6). The turquoise module in the EY tissue was significantly associated with differences in social exposure with female in *H. m. malleti* (Figure S6A, B). This module is enriched for various mRNA processes and zinc ion binding (Table S29, S30). Four modules in the antennae (turquoise, lightcyan, yellow, saddlebrown) were uniquely associated with training in *H. m. malleti* suggesting that gene networks may be important when males are associating female experience-based preferences (Figure S6D, Table S32, S33, S34, S35). GO enrichment identified 17 most specific (FDR<0.05) terms for genes in the turquoise module (Figure S6E, Table S32, S33), most notable of which was zinc ion binding, which is needed for several neural development and nervous system modulations^5^.

***Other putative “magic” trait genes between H. m. malleti and H. m. rosina:***

Genes within the loci responsible for the development of the divergent trait, and the preference for that trait are hotspots for trait-preference co-evolution. Genes associated with differences in mate preference learning that are present in these hotspot loci will therefore undergo correlated divergent selection due to linkage, thereby enabling divergence in mate preference learning as well. In butterflies, and especially in *Heliconius,* wing color and pattern are “magic” traits, that are under divergent natural selection, but also under sexual selection as these traits are used by both sexes in assortative mating^6–8^.

We found 20 known butterfly wing patterning genes, of which 8 were differentially expressed across populations in at least one of the three tissues during the control or training treatments (Table S36). Most notable of these were three different *yellow* genes that were upregulated in the brains of *H. m. malleti.* Two of these (*y*, HMEL021221g1, log2foldchange = 0.89; and *yellow-f2,* HMEL022544g1, log2foldchange = 2.42) were upregulated in the brain during the control treatment, while the third (*y*, HMEL008064g1, log2foldchange = 0.59) was upregulated in the brain during the training treatment. Not only does the *yellow* gene regulate some butterfly species’ wing colors and pattern development through melanin pathway, it also influences male courtship behavior in *Bicyclus anynana*^9^.

Two other wing patterning genes that are essential in *Heliconius* wing development are *aristaless 1*^10^ (HMEL011985g1), and *WntA*^11^ (HMEL018100g1). *Aristaless 1* acts downstream of *WntA* and *cortex* to differentiate white and yellow scale colors, whereas *WntA* expression delineates scale cell boundaries during wing development^12^. A gene adjacent to *aristaless 1* was DE in the brains between *H. m. malleti* and *H. m. rosina. Alsin* (HMEL11986g1)– which regulates synaptic transmissions– was downregulated in *H. m .malleti* brains during training conditions. A gene (HMEL002905) adjacent to *WntA* that is annotated for juvenile hormone esterase binding protein was upregulated in the brains of *H. m. malleti* males during training.

***Learning and memory DEGs on the Z sex chromosome:***

We assessed whether genes that were expressed across the three tissues were biased to any one particular chromosome in the genome. We found that genes were evenly expressed across all the 21 chromosomes in all three tissues. In the brain, gene expression ranged between 80-92% across the genome (Figure S7A), in the eye it ranged between 76-88% (Figure S7B), and in the antennae it ranged between 79-92% (Figure 7C). However, the Z sex chromosome (chromosome 21) had the highest DEGs, both by percentage, and by numbers, between *H. m. malleti* and *H. m. rosina* in all the three tissues (Figure S8 A, B, C), irrespective of their exposure with a female. This suggests that the Z sex chromosome have diverged between the two subspecies.

The sex chromosomes are hypothesized to play an important role in reproductive isolation^13^. When specifically examining genes on the Z sex chromosome (chromosome 21), we found 159 DEGs in the brain, 136 in the eyes, and 101 in the antennae during an exposure with a female (training condition). Some notable genes are *protein henna* (HMEL011020g1, log2foldchange = 2.8 in antennae) which is a part of the dopamine biosynthesis pathway and implicated in long term memory formation in *Drosophila melanogaster*^14^, and *Gamma-aminobutyric acid receptor subunit beta* (HMEL003497g3, log2foldchange = -0.37 in brain) which regulates neurotransmitter activity controlling various behaviors (aggression, learning, sleep), and over-expression of which is known to inhibit memory formation in *D. melanogaster*^15–17^.
